# European LD proxies lose tagging for 43% of GWAS leads in the African panel: a chromosome 22 analysis with SwiftLD, an 826-fold faster, byte-identical vcftools LD

**DOI:** 10.64898/2026.11.04.760946

**Authors:** Joe Christensen

## Abstract

SwiftLD is a drop-in replacement for the pairwise linkage disequilibrium (LD) modes of vcftools 0.1.17, `--hap-r2` and `--geno-r2`, which population-genetic pipelines use to compute r^2^, D and D′ between nearby variants. On three regions of 1000 Genomes chromosome 22 with 503 to 2,504 samples it wrote the same output as vcftools, byte for byte, and was 826-fold faster (geometric mean at two threads on an AVX-512 CPU). It was also 7.6-fold faster than PLINK 2, the fastest alternative, from VCF input to LD output on all five regions tested; PLINK 2 writes a different file, although its r^2^ values agree with vcftools’ within 10^−6^. One megabase with 503 samples took vcftools 29 s and SwiftLD 0.049 s, and on an ordinary AVX2 laptop SwiftLD was 102-fold faster than vcftools at 12 threads. vcftools was slow because it repeatedly re-reads genotypes from a temporary file, counts haplotypes individually and flushes every output row. I built SwiftLD with correctness-gated coding agents (Claude Opus 5.5): every change had to keep the output byte-identical to vcftools. No output byte differed in 167,749 differential tests on three machines. On chromosome 22, the best European LD proxy of a GWAS lead kept r^2^ ≥ 0.8 for 57.4% of leads in the African panel and 85.6% to 92.7% in other panels (98.1% in a European split-half control). Code and a two-command reproduction: https://github.com/joe-b-christensen/swiftld.

## Introduction

Linkage disequilibrium (LD), the non-random association of alleles at different loci, enters many population-genetic analyses: pruning markers before principal component and admixture analyses, estimating recombination and effective population size from the decay of LD with distance, and fine-mapping association signals (Slatkin 2008). vcftools (Danecek et al. 2011), cited more than 16,000 times, is a standard way to compute it. `--hap-r2` reports r^2^, D and D′ from phased haplotypes (Lewontin 1964; Hill and Robertson 1968), `--geno-r2` the squared correlation of genotype dosages, and both read a VCF file directly and apply vcftools’ site and individual filters. They are slow: one megabase of 1000 Genomes data with 503 samples and a 200-kb window takes 29 s. PLINK 1.9 and 2.0 (Chang et al. 2015) and scikit-allel (Miles et al. 2024) are faster, but they need an import step, define filters differently and print other rows, so switching to them changes downstream numbers.

SwiftLD writes the same output file as vcftools, byte for byte. On three 1000 Genomes regions it was 826-fold faster than vcftools at two threads, and on five it was 7.6-fold faster than PLINK 2 (Fig. 1), so a pipeline can switch without re-validating a single number. The waste in vcftools is simple. It keeps every site in a temporary file and, for every pair, reads the second site back one byte at a time and counts it one haplotype at a time. Decoding each site once into bit-planes and counting pairs with popcount gave the first 77.5-fold; making input and output as cheap as the counting gave the rest.

**Figure 1.**
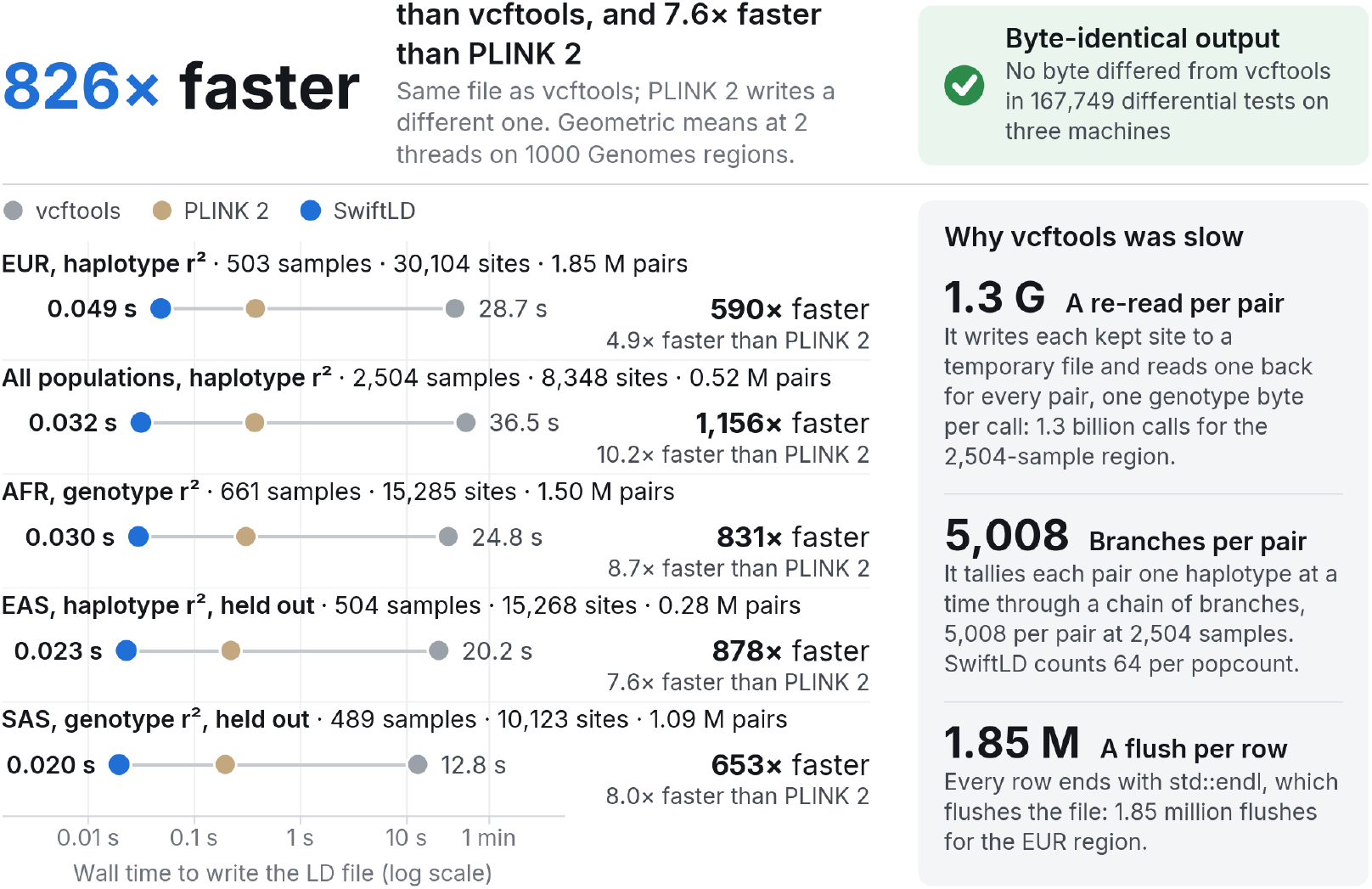
SwiftLD writes the same file as vcftools in a few hundredths of a second where vcftools takes 13 to 36 s, and is 4.9- to 10.2-fold faster than PLINK 2, which writes a different file. Wall time of vcftools 0.1.17 (grey; one thread), PLINK 2 (sand; import plus LD) and SwiftLD (blue), the last two at two threads, on five regions of 1000 Genomes chromosome 22, log scale; the bottom two regions were held out from development. AVX-512 cloud VM with 2 vCPUs. PLINK 2 ratios use SwiftLD times from the same benchmarking script (Section 4.5). Right: the three habits that made vcftools slow (Section 2.4).

LD also determines whether a marker that tags a GWAS signal in one reference panel tags it in another. I measured LD decay in all 26 populations of 1000 Genomes and tested how European proxies transfer to other panels, using chromosome 22 only.

## 2 Results

### 2.1 SwiftLD is 826-fold faster than vcftools on 1000 Genomes data

On the three 1000 Genomes development regions SwiftLD was 826-fold faster than vcftools at two threads and 402-fold faster at one (geometric means; Fig. 1, Table 1). The largest factor came on the region with all 2,504 samples, where vcftools took 36 s and SwiftLD 0.032 s (1,156-fold), because vcftools’ cost per pair grows with the number of samples (Section 2.4). On the two held-out regions, which were never used during development, SwiftLD was 878-and 653-fold faster. Across all eleven benchmark sets, which add six synthetic sets with missing and unphased calls, extra FORMAT fields, several chromosomes and 1,500 samples, it was 163-to 1,309-fold faster; the slowest is a set where 5% missing calls send every pair through the general counting path.

**Table 1.**
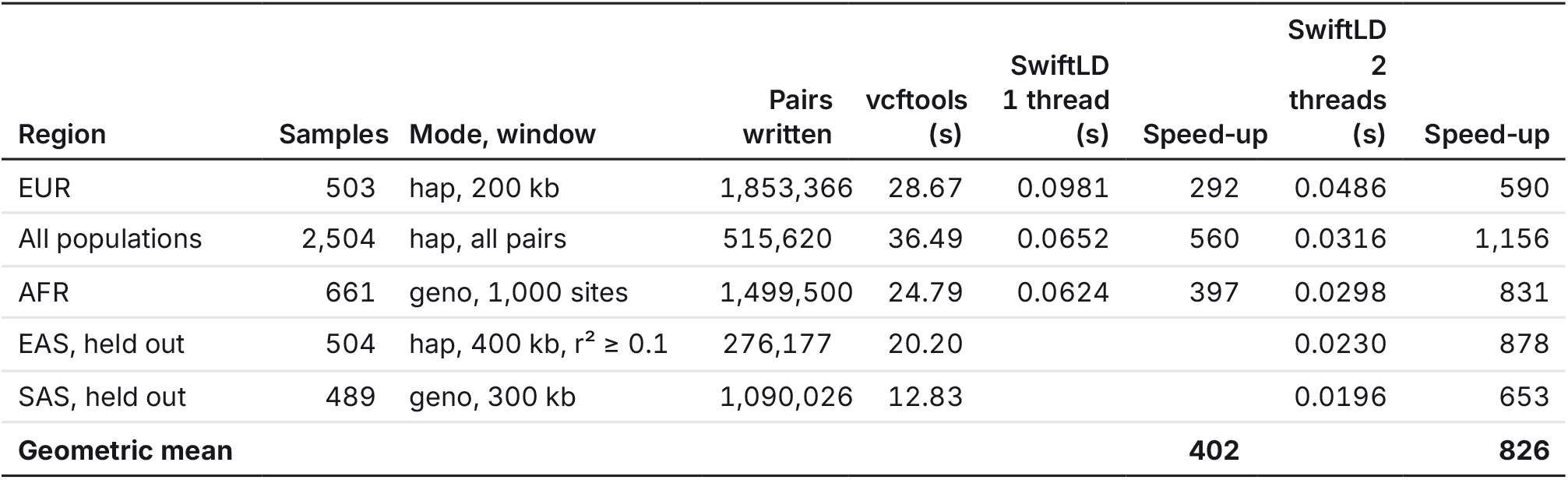
SwiftLD wrote byte-identical output on every 1000 Genomes region and was 590- to 1,156-fold faster than vcftools at two threads. AVX-512 cloud VM with 2 vCPUs; vcftools 0.1.17 best of three runs; SwiftLD at two threads, median of three evaluation runs (held-out regions, one run), each the best of up to three; at one thread, one run. Mode: haplotype r^2^ (`--hap-r2`) or genotype r^2^ (`--geno-r2`); minor allele frequency thresholds and the full options are in Supplementary Table S1. The geometric means are over the three development regions; at two threads it is the median of the three runs’ geometric means (the geometric mean of the per-region medians is 827).

The speed-up holds on other machines (Fig. 2). On a second cloud VM of the same type, with vcftools re-timed there, SwiftLD was 541-fold faster on the development regions at two threads; on that VM vcftools ran 11–17% faster and SwiftLD 8–51% slower than on the first. On a laptop without AVX-512 (AMD Ryzen 5 5600H, 12 threads), where the build selected the portable code paths, SwiftLD was 102-fold faster on the development regions and 89-fold faster on the held-out regions.

**Figure 2.**
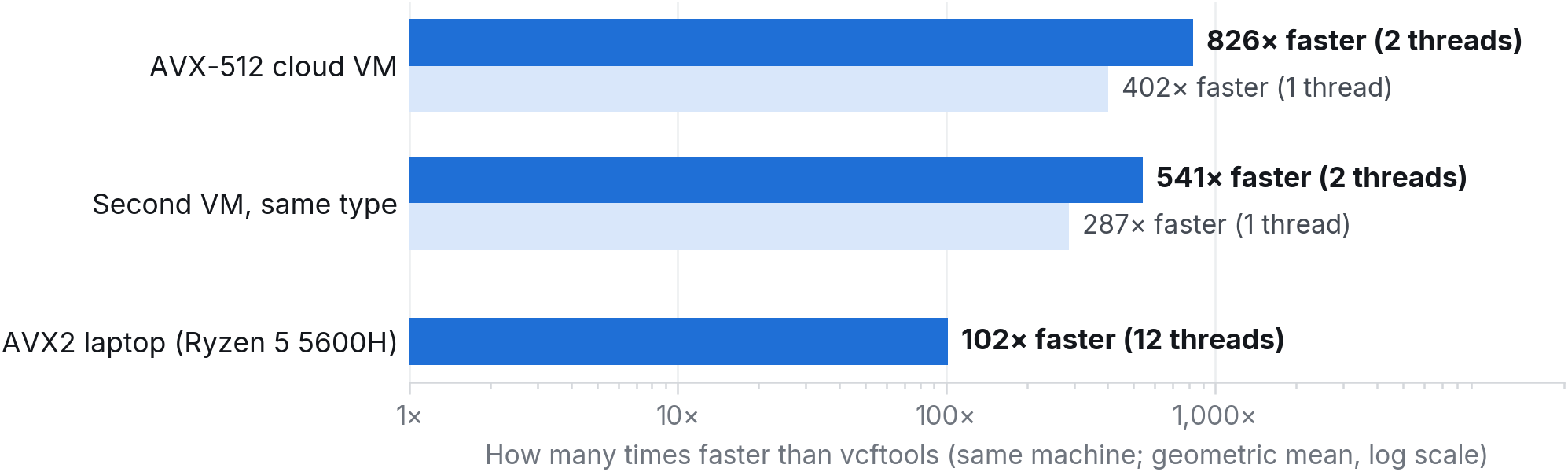
SwiftLD is faster than vcftools on every machine tested. Geometric mean over the three 1000 Genomes development regions of how many times faster SwiftLD was than vcftools 0.1.17 on the same machine: the AVX-512 VM of Table 1, a second VM of the same type with vcftools re-timed there, and an AVX2 laptop (AMD Ryzen 5 5600H) on which the build selected the portable code paths. Dark bars, all threads; light bars, one thread (not measured on the laptop); log scale. Every output was byte-identical to vcftools’.

### 2.2 The output is identical, byte for byte

No output byte differed between SwiftLD and vcftools in 167,749 differential tests (Table 2). Every code change had to pass a check that ran 300 fresh random VCF files through both programs, with random options and thread counts, and compared the output on six benchmark sets with checksums recorded for vcftools; 77 such checks add up to 23,100 random tests. Five held-out sets passed every time they were checked. On the final code, edge-case tests, among them 34-to 63-MB gzip inputs and builds without AVX-512, and separate random runs added 142,549 cases, and the full check passed on a second VM and on a laptop with a different CPU vendor. Byte identity covers row order, six-significant-digit formatting and the sign of NaN, which vcftools prints as `-nan`.

**Table 2.** No output byte differed in any comparison. Each test compared the complete output of SwiftLD and vcftools 0.1.17 on the same input, options and thread count; benchmark outputs were compared by sha256.

| Test | Runs | Result |
|---|---|---|
| Random VCF files against the live vcftools during development | 23,100 | all identical |
| Held-out sets, never used during development (5 sets × 6 checks) | 30 | all identical |
| Edge-case tests on the final code (large gzip inputs, builds without AVX-512) | 22,549 | all identical |
| Separate random VCF files on the final code | 120,000 | all identical |
| Numbers formatted by SwiftLD against glibc snprintf with %.6g, vcftools’ format | 83,882,953 | all identical |
| Fresh copy of the code: five 1000 Genomes outputs at 1, 2 and 3 threads | 15 | all identical |
| Full check on a second VM of the same type: random VCF files | 1,500 | all identical |
| Full check on an AVX2 laptop (AMD Zen 3): random VCF files | 600 | all identical |

### 2.3 SwiftLD is 7.6-fold faster than PLINK 2, which writes a different file

SwiftLD was the fastest tool on every 1000 Genomes region (Figs. 1 and 3). From VCF input to LD output it was 4.9- to 10.2-fold faster than PLINK 2 (7.6-fold on average), 5.4- to 10.9-fold faster than PLINK 1.9 and 103- to 252-fold faster than scikit-allel. Most of PLINK 2’s time goes to importing the VCF, 42–70% of its total; on data PLINK 2 had already imported, SwiftLD was still 2.2- to 4.7-fold faster than PLINK 2’s LD step alone.

**Figure 3.**
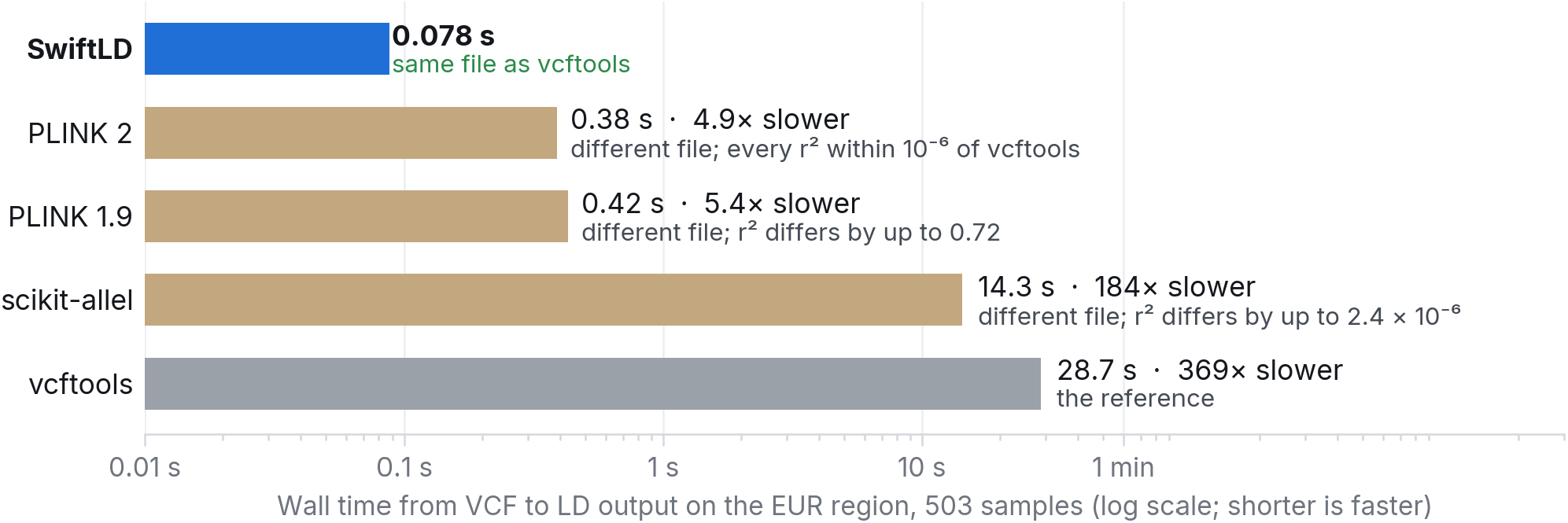
On the EUR region every other tool is slower than SwiftLD, and none of them writes vcftools’ file. Wall time from VCF input to LD output on 1000 Genomes EUR (503 samples, 1 Mb, 1.85 million pairs), log scale; PLINK times include importing the VCF and scikit-allel’s include Python start-up. PLINK at two threads on the AVX-512 VM; one script timed every tool, SwiftLD included, which is why SwiftLD’s time here is 0.078 s rather than the 0.049 s of Table 1. Agreement is with vcftools’ r^2^ on the pairs both tools reported. Supplementary Figure S1 shows all five regions; versions are in Section 4.5.

None of the three writes vcftools’ file. PLINK 2 agreed with vcftools on every shared r^2^ within 10^−6^ and scikit-allel on at least 99.96% of them, with the largest difference 4.8 × 10^−6^. PLINK 1.9’s `--r2` agreed on the genotype regions, but on phased data it uses an unphased composite estimator and differed by up to 0.79. All three print their own rows and columns, define allele frequency and missingness on their own imported data and do not print D′ as vcftools does, so a pipeline that switches to them has to re-validate its results (Supplementary Section S2).

### 2.4 vcftools re-reads and re-decodes a site for every pair

Three habits of vcftools account for most of its run time (Fig. 4).

**Figure 4.**
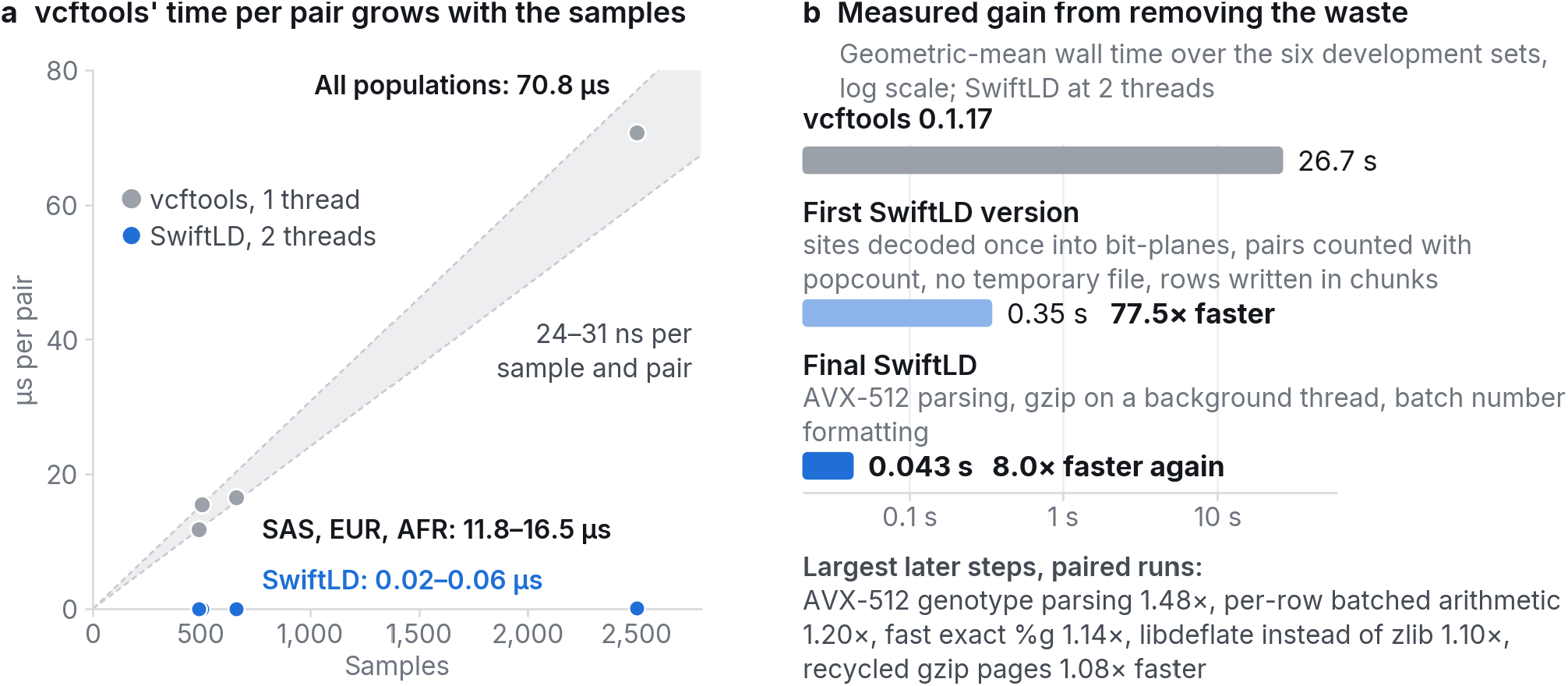
vcftools’ cost per pair grows with the number of samples, and removing that waste gave the first 77.5-fold. (a) Wall time of vcftools 0.1.17 divided by the pairs it wrote, against the number of samples, on four 1000 Genomes regions (grey), and the same for SwiftLD at two threads (blue); the shaded wedge spans 24 to 31 ns per sample and pair. The EAS region is left out because `--min-r2 0.1` hides the pairs vcftools computed but did not print. (b) Geometric-mean wall time over the six development sets of vcftools (best of three runs), the first SwiftLD version (one run) and the final SwiftLD (median of three runs), with the largest later gains, each measured in paired runs against the version before it.

#### It re-reads a site for every pair

vcftools writes every kept site to a temporary file, one byte per individual, and for each pair reads the second site back and decodes it into a vector of allele pairs, with one `ifstream::read` call per byte. On the 2,504-sample region that is 2,504 calls for each of 515,620 pairs, 1.3 billion in all.

#### It counts one haplotype at a time

It then walks the two sites haplotype by haplotype (individual by individual for `--geno-r2`) through a chain of branches and double-precision accumulators: 5,008 steps per pair at 2,504 samples.

#### It flushes after every row

Each output row ends with `std::endl`, which flushes the stream, 1.85 million times for the 1.85 million rows of the EUR region.

The first two make the cost of a pair proportional to the number of samples. vcftools’ run time divided by the pairs it wrote was 12 to 17 µs on the regions with 489 to 661 samples and 71 µs on the region with 2,504, or 24 to 31 ns per sample and pair (Fig. 4a).

SwiftLD decodes each site once into bit-planes over the haplotypes, or over the individuals for `--geno-r2`, so the counts of a pair are popcounts of ANDed 64-bit words. It then evaluates vcftools’ double-precision expressions in the same order from the exact integer counts, compiled without fused multiply-add, so every value agrees to the last bit. The first version did this and wrote rows in chunks from worker threads, strictly in order; it was 77.5-fold faster than vcftools on the six development sets (Fig. 4b). Input and output then dominated, and the next 8.0-fold came from validating and decoding regular genotype lines 64 bytes at a time with AVX-512, decompressing gzip with libdeflate (Biggers 2026) on a background thread that the parser follows, an AVX-512 formatter for vcftools’ `%.6g` numbers that sends any value near a rounding tie to an exact path, and recycling the memory pages the parser had passed. Because rows are written in order, the output does not depend on the thread count.

### 2.5 European proxy retention on chromosome 22 ranges from 57.4% to 92.7% across target panels

Half-decay distances ranged from 4.3 to 11.7 kb across 26 populations sampled at 61 individuals each: 4.3 to 4.8 kb in the seven African populations and 8.0 to 11.7 kb in the other 19. LD was computed at a fixed random 5% of biallelic SNP positions shared across populations, retaining within-sample MAF ≥ 0.05 (Fig. 5; Supplementary Table S4). The long-distance mean at 400–500 kb ranged from 0.010 to 0.018. The intervals describe variation among blocks of this chromosome.

**Figure 5.**
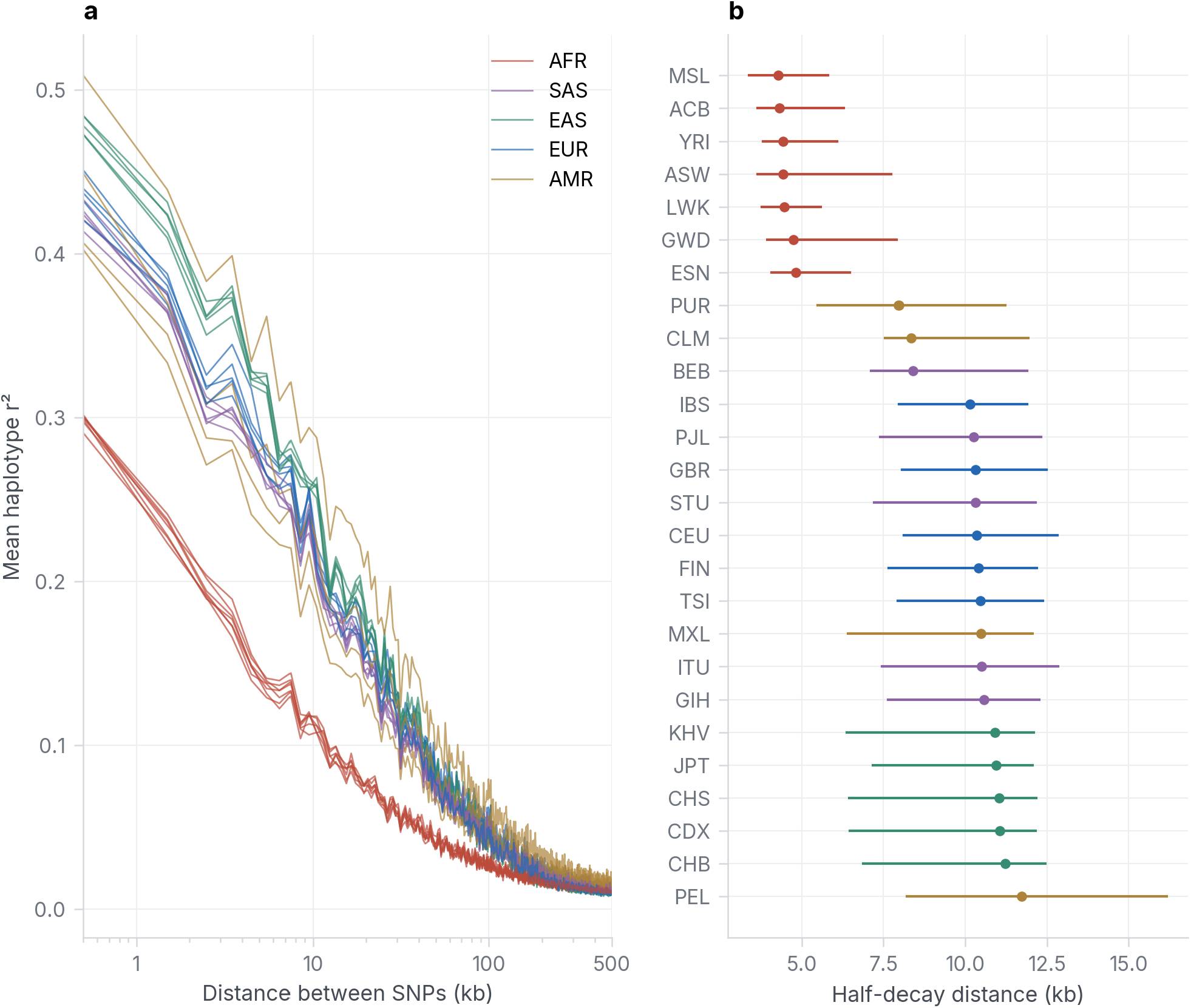
LD decay differs among the 26 population samples on chromosome 22. (a) Mean haplotype r^2^ in 1-kb distance bins, coloured by the Phase 3 super-population labels; log distance axis. (b) Distance at which mean r^2^ first falls to half the first-bin mean, sorted by the estimate. Points use all pairs of eligible selected SNPs; bars are percentile 95% intervals from 1,000 resamples of 1-Mb blocks assigned by the first SNP. Each population contributes 61 individuals.

The dated GWAS Catalog supplied 4,043 unique lead rsIDs on chromosome 22 with EUR MAF ≥ 0.05, of which 3,544 had at least one European proxy at r^2^ ≥ 0.8 within 500 kb; 499 had none. The best European proxy retained r^2^ ≥ 0.8 at these rates: African (AFR) 57.4%, median r^2^ 0.88; American (AMR) 92.7%, median r^2^ 0.98; East Asian (EAS) 85.6%, median r^2^ 0.99; South Asian (SAS) 92.1%, median r^2^ 0.99 (Fig. 6). The split-half EUR control retained 98.1% of 3,555 eligible pairs, with median r^2^ 0.99. Its proxy selection used the discovery half alone. The target comparisons and control therefore have different proxy sets and denominators.

**Figure 6.**
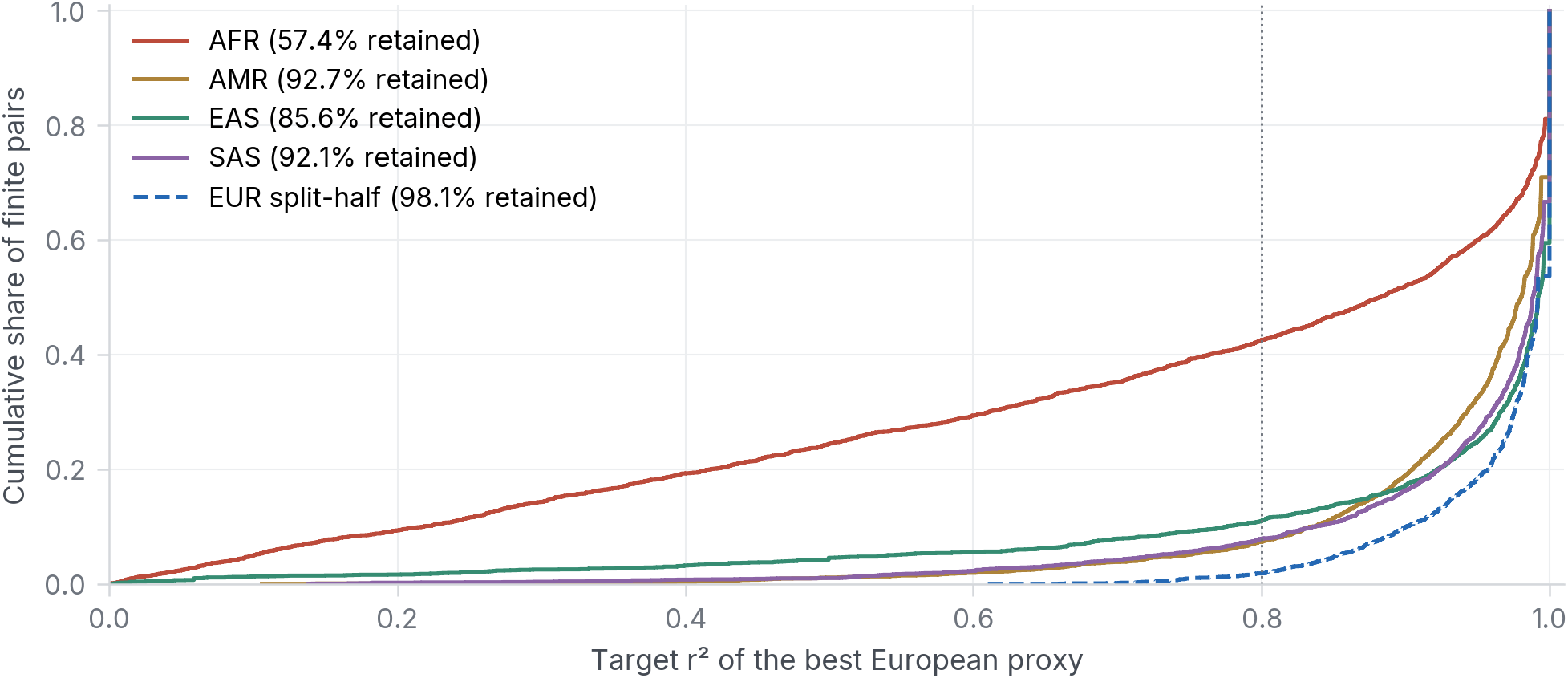
The r^2^ of the best European proxy differs across target panels on chromosome 22. Cumulative distributions include finite lead–proxy r^2^ values; labels give the percentage retained at r^2^ ≥ 0.8 among all leads with a discovery proxy, including undefined target pairs as failures. The dotted line marks 0.8. The dashed curve is an independently selected EUR split-half control. AFR, AMR, EAS and SAS use all Phase 3 individuals in each panel.

The per-population results, leads below 1% target MAF, undefined pairs and sensitivity analyses at r^2^ thresholds 0.6 and 0.9 and a 250-kb window are in Supplementary Section S3. Byte comparisons with vcftools matched exactly on a 20–21-Mb ASW window (180,427 bytes) and on a 30–31-Mb window of the European discovery input (61,996 pairs). After input acquisition, preparation, LD computation, checks and figure generation took 0.47 h on the 12-thread laptop. European proxy discovery took SwiftLD 3.1 s; vcftools would take an estimated 21 min, extrapolating its measured 11.9 s on the 30–31-Mb window by the number of SNP pairs within 500 kb. The 26 decay scans, on their 5% sample of positions, would take vcftools an estimated 3.3 min. The new analysis covers one chromosome.

## 3 Discussion

SwiftLD’s margin over vcftools comes from removing work the answer does not need, not from approximation. vcftools’ arithmetic is deterministic and its inputs are small integers, so its own expressions, fed exact counts, reproduce every bit. The margin over PLINK 2 is smaller, 4.9- to 10.2-fold, because PLINK 2 already counts pairs with bit operations; most of its remaining time goes to importing the VCF.

Byte identity is a stronger promise than agreement on a test set, and it is the promise that lets a user switch without re-validation. Testing every change against vcftools made it checkable while the code changed. It forced SwiftLD to reproduce vcftools’ filter order, its window scan, the sign of its NaN and the defects that shape its output, and it forced an exact fallback behind every fast path. The protection held on data the tests had never seen: on held-out regions, on a second VM and on a laptop with a different CPU vendor and no AVX-512. The same approach carries over to any tool whose output can be compared byte for byte.

The chromosome 22 analysis gives a practical reason to compute LD in the target reference panel. On chromosome 22, the best European LD proxy retained r^2^ ≥ 0.8 for 57.4% of GWAS lead SNPs in AFR and 92.7% in AMR, compared with 98.1% in a European split-half control. This comparison concerns marker tagging in these panels; it does not measure transfer of causal effects or prediction accuracy.

### Limitations

The headline was 826-fold on the VM used for development, where single runs varied by up to 20%, and 541-fold on a second VM of the same type. It uses two threads, all CPUs of the VM, against single-threaded vcftools, and scaling beyond two threads was not measured. Timed inputs were single-member gzip regions of 0.25 to 1 Mb; the new chromosome 22 application used uncompressed VCF input, while BGZF input, which SwiftLD reads through a slower non-streaming path, was not timed. Identity with vcftools rests on differential tests over the 18 supported options, not on proof. As byte identity requires, SwiftLD copies the vcftools defects that shape the output, such as genotypes reused from the previous record on short lines (vcftools issue #234), and it exits with an error on corrupt gzip input where vcftools hangs (#235).

A SwiftLD output file is a vcftools output file, so every pruning list, decay curve and fine-mapping input built from it stays as it was. The 29-second vcftools run on one megabase of EUR haplotypes takes SwiftLD a twentieth of a second, and on an ordinary laptop without AVX-512 SwiftLD was 71- to 117-fold faster than vcftools on each of the five regions.

## 4 Methods

### 4.1 Target

I took the pairwise LD modes of vcftools 0.1.17 (commit 1c53c3c) as the target. SwiftLD accepts 18 vcftools options in any combination and order (`--vcf`, `--gzvcf`, `--out`, `--stdout`; `--ld-window`, `--ld-window-min`, `--ld-window-bp`, `--ld-window-bp-min`, `--min-r2`; `--chr`, `--maf`, `--max-missing`, `--remove-filtered-all`, `--remove-indels`, `--thin`, `--phased`; `--keep`, `--remove`), writes output identical byte for byte to vcftools’ at any thread count (SWIFTLD_THREADS, default all CPUs) and exits non-zero wherever vcftools fails. The reference binary carried one upstream fix, for the temporary-file name buffer (vcftools issue #231), without which every LD run aborts on Ubuntu. SwiftLD vendors libdeflate 1.26 with three small patches. I developed SwiftLD with correctness-gated coding agents (Claude Opus 5.5, Anthropic), following SwiftNJ (Christensen 2026): every code change had to pass the correctness test in Section 4.2.

### 4.2 Correctness testing

Each test ran SwiftLD and the live vcftools binary on the same random input: a small adversarial VCF file with 1 to 65 samples and up to 120 records, mixing phased, unphased, haploid and triploid calls, missing data, multi-allelic, indel and `ALT=.` records, unsorted positions, interleaved chromosomes and gzip input, run with random options and 1 to 7 threads. The output had to match byte for byte, and a failing vcftools run had to be matched by a failing SwiftLD run. Each check ran 300 such tests from a fresh seed and compared the output on each of the six development sets with the checksum recorded for vcftools. The five held-out sets were checked six times and never used to choose between versions.

### 4.3 Data

The real data are five regions of the 1000 Genomes Phase 3 release of chromosome 22 (GRCh37; 2,504 samples phased with SHAPEIT2; 1000 Genomes Project Consortium 2015), pinned by sha256, with every record kept. The three development regions are EUR (503 samples, 1 Mb), all populations (2,504 samples, 0.25 Mb) and AFR (661 samples, 0.5 Mb, phase removed for `--geno-r2`); the two held-out regions are EAS (504 samples, 0.5 Mb) and SAS (489 samples, 0.3 Mb, phase removed). Three synthetic development sets and three synthetic held-out sets from a haplotype-mosaic generator add all-pairs output, missing and unphased calls, extra FORMAT fields, multi-allelic records, several chromosomes and 1,500 samples (Supplementary Table S1).

### 4.4 Timing

Times are wall times of the whole command, including reading, decompressing, computing and writing the output to a file. Headline timings come from a cloud VM with 2 vCPUs (Intel Xeon at 2.1 GHz, AVX-512, 7.8 GB RAM, g++ 13.3.0). vcftools ran three times per set and the best time counts.

SwiftLD’s two-thread times are the median of three evaluation runs on the final code, each per-set time the best of up to three runs; the held-out sets and the one-thread times come from one evaluation run each. The headline is the median of the three runs’ geometric means over the development regions, and every speed-up is relative to vcftools on the same machine. The whole protocol was repeated on a second VM of the same type, with vcftools re-timed there, and on a laptop (AMD Ryzen 5 5600H, Zen 3, 6 cores and 12 threads, AVX2 without AVX-512, WSL2 Ubuntu 24.04, gcc 13.3) at 12 threads.

### 4.5 Baselines

PLINK 2.0 and PLINK 1.9 (Chang et al. 2015; plink-ng commit 03bc669) and scikit-allel 1.3.13 (Miles et al. 2024) ran on the five 1000 Genomes regions on the same VM, with windows and allele frequency thresholds matching vcftools’: PLINK 2 `--r2-phased` or `--r2-unphased` and PLINK 1.9 `--r2`, both at two threads after importing the VCF, and scikit-allel `rogers_huff_r` (Rogers and Huff 2009) on haplotypes or dosages. One script timed every tool, SwiftLD included, as the best of two runs for runs under 5 s; it timed SwiftLD at 0.025 to 0.078 s, against 0.020 to 0.049 s in Table 1, and every ratio to another tool uses SwiftLD’s time from that script. Agreement of r^2^ with vcftools was measured on the pairs both tools reported (Supplementary Section S2).

### 4.6 Chromosome 22 application

I used the Phase 3 GRCh37 chromosome 22 v5b VCF and sample panel (1000 Genomes Project Consortium 2015), and the GWAS Catalog associations release dated 2026-10-22 (Cerezo et al. 2025), downloaded on 4 November 2026. Input URLs, byte sizes and SHA256 checksums are in application/results/inputs.json. The chromosome contained 1,103,547 records, including 1,055,454 biallelic SNPs. The v5b VCF lacked rsIDs. Catalog chromosome 22 rsIDs were resolved at https://grch37.rest.ensembl.org/variation/homo_sapiens; uniquely mapped GRCh37 chromosome 22 SNPs were matched to v5b by position and compatible alleles. Request lists, response file names and SHA256 checksums are recorded in application/results/rsid_mapping_inputs.json.

For LD decay I sampled 61 individuals without replacement per population, the size of the smallest panel (ASW), using seed 20261104. A fixed random 5% of unique biallelic SNP positions was drawn once with seed 20261104 and shared across populations; each population then retained within-sample MAF ≥ 0.05. This was the predefined fallback after the full-site run proved too slow for the available runtime. SwiftLD computed phased haplotype r^2^ within 500,000 bp at 12 threads. A streaming C++ binner accumulated pair counts and r^2^ sums in distance bins (0,1,000], (1,000,2,000], and so on. Half-decay was the first crossing of half the first-bin mean, interpolated between bin centres. I resampled occupied 1-Mb physical blocks 1,000 times, assigning each pair to its first SNP, and took percentile 95% intervals. Summaries at 10, 50 and 100 kb refer to bins ending at those distances; the baseline pools all pairs at 400–500 kb.

For proxy transfer I kept Catalog rows with p ≤ 5 × 10^−8^ whose chromosome field was 22 and SNPS field contained one rsID, deduplicated rsIDs, matched biallelic chromosome 22 SNPs, and required EUR MAF ≥ 0.05. All 503 EUR individuals defined the discovery panel. Proxies excluded the lead, lay within 500 kb and had r^2^ ≥ 0.8. The best proxy maximised EUR r^2^, breaking ties by distance then position.

Thresholds and ranking used SwiftLD’s six-significant-digit output, matching vcftools. Targets used all individuals in AFR, AMR, EAS and SAS and each of the 26 populations. Retention used all leads with a discovery proxy as its denominator; undefined target r^2^ counted as failure, and medians used finite values. I also recorded the fraction of each lead’s discovery proxies retained and leads with target MAF < 0.01. Seed 20261104 split EUR into 251 discovery and 252 validation individuals. Sensitivities repeated discovery and transfer at thresholds 0.6 and 0.9 and with a 250-kb window. An exact allele-frequency bound excluded discovery SNPs mathematically unable to reach r^2^ ≥ 0.6 with any eligible lead; it did not remove any possible proxy. All LD output was piped into summaries or lead-pair selection. Hand-computed examples checked bin sums, boundaries and proxy selection; a fixed 1-Mb subsampled ASW window checked complete output byte identity against vcftools.

### 4.7 Use of AI

The software and this text were made in collaboration with Claude Opus 5.5 (Anthropic) from my spoken instructions. I chose the problem, gave the instructions, reviewed the results and approved the final text.

## Data and code availability

The code is available at https://github.com/joe-b-christensen/swiftld and archived on Zenodo (DOI: 10.5281/zenodo.23963404). SwiftLD is free software under the GNU Lesser General Public License v3.0, the licence of vcftools, from whose behaviour it derives; it vendors libdeflate 1.26 (MIT licence) with three small patches. The repository contains the source code, its build script, the scripts that check the output against vcftools and time both programs (prepare.py, evaluate.py, gen_data.py) and the five 1000 Genomes inputs.

The synthetic datasets are regenerated deterministically by gen_data.py. The real inputs are subsets of the 1000 Genomes Project Phase 3 release (1000 Genomes Project Consortium 2015), file ALL.chr22.phase3_shapeit2_mvncall_integrated_v5b.20130502.genotypes.vcf.gz and sample panel integrated_call_samples_v3.20130502.ALL.panel from http://ftp.1000genomes.ebi.ac.uk/vol1/ftp/release/20130502/, retrieved on 11 October 2026. These data are openly available without restriction under the data reuse policy of the International Genome Sample Resource (https://www.internationalgenome.org). The five subsets (4.7 MB) are in the repository’s `data/` folder with their sha256 checksums, together with the scripts that recreate them from the release file.

The chromosome 22 analysis scripts are in the repository’s application/scripts/ directory. The full chromosome VCF and dated GWAS Catalog associations remain available from the source URLs recorded with their checksums in application/results/inputs.json.

## Competing interests

None declared.

## Funding

None.

## Notes

https://github.com/joe-b-christensen/swiftld

### Competing Interest Statement

The authors have declared no competing interest.

