## Supplementary Material for "European LD proxies lose tagging for 43% of GWAS leads in the African panel: a chromosome 22 analysis with SwiftLD, an 826-fold faster, byte-identical vcftools LD"

**European LD proxies lose tagging for 43% of GWAS leads in the African panel: a chromosome 22 analysis with SwiftLD, an 826-fold faster, byte-identical vcftools LD**
Joe Christensen (ORCID 0009-0007-0908-1299). Translational Neuropsychiatry Unit (TNU), Department of Clinical Medicine, Aarhus University, Denmark.

### S1 Datasets

The synthetic sets are generated by gen_data.py from fixed seeds; the real regions are described in main-text Section 4.3 and in the data availability statement.

**Table S1.** The eleven benchmark sets. Sites are VCF records in the file, before vcftools' filters. The real regions of chromosome 22 (GRCh37) are 17.00–18.00 Mb (EUR), 18.00–18.25 Mb (all), 18.25–18.75 Mb (AFR), 19.00–19.50 Mb (EAS) and 19.50–19.80 Mb (SAS); every record in a region is kept, and the genotype sets had every | replaced by / to make unphased input for `--geno-r2`. The synthetic sets come from a haplotype mosaic (founder haplotypes with Beta(0.35, 0.35) allele frequencies, switching founder with probability $1 - \exp(-\rho \times \text{gap})$) with missing data, unphased calls, extra FORMAT fields, multi-allelic records, indels, several chromosomes and gzip input layered on top.

| Set | Region / role | Samples | Sites | Options |
|---|---|---|---|---|
| r_eur_hap | EUR, development | 503 | 30,104 | `--hap-r2 --maf 0.05 --ld-window-bp 200000` |
| r_all_hap | all populations, development | 2,504 | 8,348 | `--hap-r2 --maf 0.05` |
| r_afr_geno | AFR, development | 661 | 15,285 | `--geno-r2 --maf 0.05 --ld-window 1000` |
| s_hap_full | synthetic, development | 150 | 2,500 | `--hap-r2` |
| s_geno_full | synthetic, development | 250 | 2,500 | `--geno-r2` |
| s_hap_multichr | synthetic, development | 200 | 24,000 | `--hap-r2 --ld-window-bp 150000` |
| rh_eas_hap | EAS, held out | 504 | 15,268 | `--hap-r2 --min-r2 0.1 --ld-window-bp 400000 --maf 0.02` |
| rh_sas_geno | SAS, held out | 489 | 10,123 | `--geno-r2 --ld-window-bp 300000 --maf 0.01` |
| h_many_samples | synthetic, held out | 1,500 | 900 | `--geno-r2` |
| h_hap_chr | synthetic, held out | 120 | 9,000 | `--hap-r2 --chr chr2 --ld-window 2000 --ld-window-min 3` |
| h_dense_geno | synthetic, held out | 60 | 6,000 | `--geno-r2 --min-r2 0.05` |

### S2 Other tools

**Table S2.** Wall times in seconds on the AVX-512 VM, one script for every tool, best of two runs for runs under 5 s; vcftools is the reference time (best of three). PLINK at two threads; scikit-allel includes Python start-up and import. PLINK 2 over SwiftLD: 4.9, 10.2, 8.7, 7.6 and 8.0-fold (geometric mean 7.6); PLINK 1.9 over SwiftLD: 5.4 to 10.9-fold; scikit-allel over SwiftLD: 103 to 252-fold. PLINK 2's LD step alone over SwiftLD: 2.2 to 4.7-fold.

| Set | vcftools | SwiftLD 2 threads | PLINK 2 import | PLINK 2 LD | PLINK 2 total | PLINK 1.9 total | scikit-allel |
|---|---|---|---|---|---|---|---|
| r_eur_hap | 28.66 | 0.0777 | 0.2057 | 0.1739 | 0.3796 | 0.4192 | 14.30 |
| r_all_hap | 36.49 | 0.0367 | 0.2593 | 0.1134 | 0.3727 | 0.4006 | 9.259 |
| r_afr_geno | 24.79 | 0.0355 | 0.1430 | 0.1647 | 0.3078 | 0.3272 | 5.175 |
| rh_eas_hap | 20.20 | 0.0294 | 0.1150 | 0.1071 | 0.2221 | 0.2259 | 5.408 |
| rh_sas_geno | 12.83 | 0.0246 | 0.0820 | 0.1147 | 0.1967 | 0.1974 | 2.538 |

**Table S3.** Agreement with vcftools' $r^2$ on the position pairs both tools reported, ranges over the five 1000 Genomes regions.

| Tool and mode | Sets | Shared pairs per set | Largest abs. difference | Fraction within $10^{-6}$ |
|---|---|---|---|---|
| PLINK 2 `--r2-phased` / `--r2-unphased` | 5 | 276,017 to 1,850,551 | 0 to $1.0 \times 10^{-6}$ | 1.0000 |
| PLINK 1.9 `--r2`, genotype sets | 2 | 1,090,026 to 1,497,637 | 0 to $1.7 \times 10^{-29}$ | 1.0000 |
| PLINK 1.9 `--r2`, haplotype sets | 3 | 267,827 to 1,850,551 | 0.65 to 0.79 | 0.0025 to 0.0588 |
| scikit-allel `rogers_huff_r` | 5 | 276,017 to 1,850,551 | $1.6 \times 10^{-6}$ to $4.8 \times 10^{-6}$ | 0.9996 to 0.9999 |

PLINK 1.9 `--r2` uses an unphased composite estimator even on phased data, hence the large differences on the haplotype sets; differences of about $10^{-6}$ reflect vcftools' six printed digits. Commands: PLINK 2 `--vcf in.vcf.gz --max-alleles 2 --make-pgen`, then `--r2-phased` or `--r2-unphased` with `--maf` and `--ld-window-kb` or `--ld-window` matching the vcftools window and `--threads 2`; PLINK 1.9 `--vcf in.vcf.gz --biallelic-only strict --make-bed`, then `--r2` with the same window; scikit-allel `allel.read_vcf`, then `allel.rogers_huff_r` on haplotypes or dosages, squared. Versions: PLINK v2.0.0-b.1 and v1.9.1 built from plink-ng commit 03bc669; scikit-allel 1.3.13.

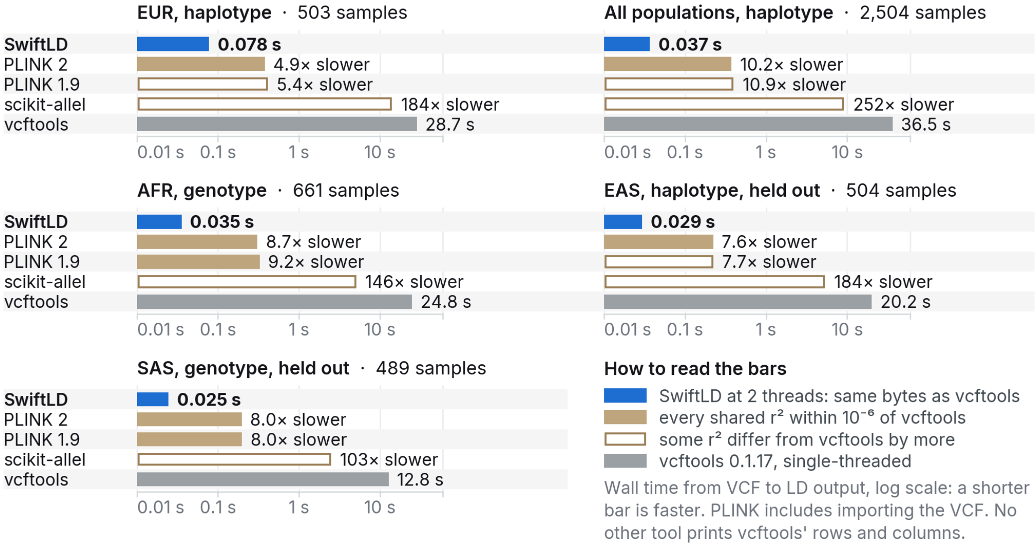


**Figure S1.** SwiftLD is the fastest tool on every 1000 Genomes region. Wall time from VCF to LD output on each region, log scale from 0.01 s; labels say how many times slower than SwiftLD each tool was, or give the time. PLINK times include importing the VCF; scikit-allel's include Python start-up. Solid bars: every $r^2$ both tools reported agreed with vcftools within $10^{-6}$; open bars: some differed by more. One script timed every tool on the AVX-512 VM, PLINK at two threads (Table S2).

### S3 Chromosome 22 application

The Catalog filters retained 1,026,666 significant association rows from 1,203,840 rows; 337,057 unique individual rsIDs, 5,832 Catalog chromosome 22 candidates, 5,436 unique GRCh37 chromosome 22 SNP mappings and 5,298 biallelic VCF matches and 4,043 EUR-common leads remained. Multi-SNP entries were excluded. The source URLs and checksums are in application/results/inputs.json.

**Table S4.** Chromosome 22 LD decay at a shared random 5% of SNP positions, equal sample sizes. Intervals resample 1-Mb first-SNP blocks.

| Population | Panel | N | Half decay, kb (95% interval) | $r^2$ at 10 kb | 50 kb | 100 kb | 400–500 kb |
|---|---|---|---|---|---|---|---|
| ACB | AFR | 61 | 4.32 (3.65–6.28) | 0.113 | 0.037 | 0.025 | 0.011 |
| ASW | AFR | 61 | 4.43 (3.65–7.74) | 0.118 | 0.036 | 0.028 | 0.013 |
| BEB | SAS | 61 | 8.39 (7.12–11.89) | 0.239 | 0.064 | 0.043 | 0.011 |
| CDX | EAS | 61 | 11.05 (6.48–12.15) | 0.256 | 0.077 | 0.050 | 0.011 |
| CEU | EUR | 61 | 10.34 (8.11–12.82) | 0.252 | 0.081 | 0.054 | 0.012 |
| CHB | EAS | 61 | 11.22 (6.89–12.45) | 0.262 | 0.073 | 0.049 | 0.011 |
| CHS | EAS | 61 | 11.04 (6.46–12.17) | 0.255 | 0.082 | 0.043 | 0.010 |
| CLM | AMR | 61 | 8.34 (7.56–11.92) | 0.218 | 0.067 | 0.049 | 0.015 |
| ESN | AFR | 61 | 4.81 (4.07–6.47) | 0.120 | 0.045 | 0.032 | 0.012 |
| FIN | EUR | 61 | 10.40 (7.66–12.19) | 0.256 | 0.086 | 0.062 | 0.014 |
| GBR | EUR | 61 | 10.31 (8.06–12.48) | 0.257 | 0.082 | 0.056 | 0.012 |
| GIH | SAS | 61 | 10.57 (7.64–12.27) | 0.247 | 0.071 | 0.050 | 0.013 |
| GWD | AFR | 61 | 4.74 (3.95–7.90) | 0.118 | 0.040 | 0.030 | 0.011 |
| IBS | EUR | 61 | 10.15 (7.98–11.89) | 0.240 | 0.080 | 0.055 | 0.011 |
| ITU | SAS | 61 | 10.49 (7.46–12.84) | 0.229 | 0.070 | 0.048 | 0.011 |
| JPT | EAS | 61 | 10.95 (7.18–12.05) | 0.258 | 0.080 | 0.048 | 0.010 |
| KHV | EAS | 61 | 10.90 (6.38–12.10) | 0.260 | 0.076 | 0.047 | 0.011 |
| LWK | AFR | 61 | 4.45 (3.77–5.57) | 0.119 | 0.039 | 0.028 | 0.012 |
| MSL | AFR | 61 | 4.28 (3.38–5.81) | 0.106 | 0.038 | 0.027 | 0.011 |
| MXL | AMR | 61 | 10.48 (6.41–12.05) | 0.241 | 0.072 | 0.057 | 0.017 |
| PEL | AMR | 61 | 11.73 (8.21–16.17) | 0.294 | 0.097 | 0.079 | 0.018 |
| PJL | SAS | 61 | 10.26 (7.41–12.31) | 0.237 | 0.070 | 0.047 | 0.011 |
| PUR | AMR | 61 | 7.96 (5.48–11.22) | 0.198 | 0.065 | 0.045 | 0.015 |
| STU | SAS | 61 | 10.31 (7.22–12.16) | 0.240 | 0.067 | 0.050 | 0.011 |
| TSI | EUR | 61 | 10.46 (7.94–12.36) | 0.244 | 0.077 | 0.050 | 0.010 |
| YRI | AFR | 61 | 4.43 (3.82–6.09) | 0.118 | 0.039 | 0.030 | 0.011 |

**Table S5.** Primary proxy transfer in all target panels. Lead MAF counts use leads with a discovery proxy. EUR_B is the independent split-half validation control. Undefined target pairs are included as failures in retention percentages.

| Target | N | Leads with proxy | Retained, % | Median r² | Mean proxy fraction retained | Lead MAF <1% | Undefined pairs |
|---|---|---|---|---|---|---|---|
| AFR | 661 | 3544 | 57.42 | 0.882 | 0.299 | 229 | 0 |
| AMR | 347 | 3544 | 92.66 | 0.980 | 0.745 | 0 | 0 |
| EAS | 504 | 3544 | 85.58 | 0.992 | 0.658 | 404 | 134 |
| SAS | 489 | 3544 | 92.10 | 0.988 | 0.712 | 62 | 0 |
| EUR_B | 252 | 3555 | 98.14 | 0.992 | 0.907 | 0 | 0 |
| ACB | 96 | 3544 | 59.48 | 0.912 | 0.327 | 58 | 30 |
| ASW | 61 | 3544 | 63.29 | 0.923 | 0.367 | 100 | 41 |
| BEB | 86 | 3544 | 90.60 | 1.000 | 0.684 | 65 | 17 |
| CDX | 93 | 3544 | 80.30 | 1.000 | 0.612 | 404 | 360 |
| CEU | 99 | 3544 | 98.73 | 1.000 | 0.929 | 0 | 0 |
| CHB | 103 | 3544 | 81.80 | 1.000 | 0.633 | 452 | 325 |
| CHS | 105 | 3544 | 81.09 | 1.000 | 0.627 | 455 | 353 |
| CLM | 94 | 3544 | 91.70 | 0.979 | 0.737 | 0 | 0 |
| ESN | 99 | 3544 | 51.55 | 0.896 | 0.262 | 418 | 409 |
| FIN | 99 | 3544 | 97.15 | 1.000 | 0.885 | 5 | 5 |
| GBR | 91 | 3544 | 98.22 | 1.000 | 0.919 | 0 | 0 |
| GIH | 103 | 3544 | 90.83 | 1.000 | 0.717 | 100 | 43 |
| GWD | 113 | 3544 | 54.20 | 0.896 | 0.283 | 380 | 232 |
| IBS | 107 | 3544 | 98.25 | 1.000 | 0.892 | 1 | 0 |
| ITU | 102 | 3544 | 90.27 | 1.000 | 0.692 | 91 | 45 |
| JPT | 104 | 3544 | 79.66 | 1.000 | 0.619 | 464 | 408 |
| KHV | 99 | 3544 | 83.58 | 1.000 | 0.635 | 322 | 197 |
| LWK | 99 | 3544 | 53.08 | 0.874 | 0.269 | 271 | 206 |
| MSL | 85 | 3544 | 50.25 | 0.880 | 0.249 | 375 | 351 |
| MXL | 64 | 3544 | 92.55 | 1.000 | 0.774 | 31 | 10 |
| PEL | 85 | 3544 | 91.14 | 1.000 | 0.765 | 66 | 16 |
| PJL | 96 | 3544 | 92.27 | 1.000 | 0.722 | 17 | 11 |
| PUR | 104 | 3544 | 88.63 | 0.980 | 0.662 | 3 | 0 |
| STU | 102 | 3544 | 90.66 | 1.000 | 0.689 | 89 | 21 |
| TSI | 107 | 3544 | 98.14 | 1.000 | 0.901 | 1 | 0 |
| YRI | 108 | 3544 | 51.78 | 0.892 | 0.271 | 452 | 388 |

**Table S6.** Threshold-specific discovery and transfer sensitivities and the 250-kb window. Each row selects its own best proxy and therefore has its own denominator.

| Target | Discovery/transfer $r^2$ | Window, kb | Leads with proxy | Retained, % | Median $r^2$ |
|---|---|---|---|---|---|
| AFR | 0.8 | 500 | 3544 | 57.42 | 0.882 |
| AFR | 0.6 | 500 | 3839 | 67.23 | 0.845 |
| AFR | 0.9 | 500 | 3247 | 51.00 | 0.910 |
| AFR | 0.8 | 250 | 3540 | 57.43 | 0.882 |
| AMR | 0.8 | 500 | 3544 | 92.66 | 0.980 |
| AMR | 0.6 | 500 | 3839 | 95.60 | 0.975 |
| AMR | 0.9 | 500 | 3247 | 86.36 | 0.985 |
| AMR | 0.8 | 250 | 3540 | 92.66 | 0.980 |
| EAS | 0.8 | 500 | 3544 | 85.58 | 0.992 |
| EAS | 0.6 | 500 | 3839 | 87.57 | 0.991 |
| EAS | 0.9 | 500 | 3247 | 83.65 | 0.995 |
| EAS | 0.8 | 250 | 3540 | 85.68 | 0.992 |
| SAS | 0.8 | 500 | 3544 | 92.10 | 0.988 |
| SAS | 0.6 | 500 | 3839 | 94.61 | 0.986 |
| SAS | 0.9 | 500 | 3247 | 88.54 | 0.991 |
| SAS | 0.8 | 250 | 3540 | 92.18 | 0.989 |
| EUR_B | 0.8 | 500 | 3555 | 98.14 | 0.992 |
| EUR_B | 0.6 | 500 | 3846 | 99.22 | 0.991 |
| EUR_B | 0.9 | 500 | 3256 | 97.02 | 1.000 |
| EUR_B | 0.8 | 250 | 3552 | 98.17 | 0.992 |

The binner and proxy hand checks passed. SwiftLD and vcftools wrote the same 180,427 bytes on the 20–21-Mb ASW window; SHA256 7be19a3c95bef532d8b271d648b4de1c61fc7d482cede82bf334c018ad478970. Their wall times were 0.044 and 0.072 s, respectively. Complete command timings are in application/results/timings.jsonl.
